# Unsupervised tensor decomposition-based method to extract candidate transcription factors as histone modification bookmarks in post-mitotic transcriptional reactivation

**DOI:** 10.1101/2020.09.23.309633

**Authors:** Y-h. Taguchi, Turki Turki

## Abstract

The histone group added to a gene sequence must be removed during mitosis to halt transcription during the DNA replication stage of the cell cycle. However, the detailed mechanism of this transcription regulation remains unclear. In particular, it is not realistic to reconstruct all appropriate histone modifications throughout the genome from scratch after mitosis. Thus, it is reasonable to assume that there might be a type of “bookmark” that retains the positions of histone modifications, which can be readily restored after mitosis. We developed a novel computational approach comprising tensor decomposition (TD)-based unsupervised feature extraction (FE) to identify transcription factors (TFs) that bind to genes associated with reactivated histone modifications as candidate histone bookmarks. To the best of our knowledge, this is the first application of TD-based unsupervised FE to the cell division context and phases pertaining to the cell cycle in general. The candidate TFs identified with this approach were functionally related to cell division, suggesting the suitability of this method and the potential of the identified TFs as bookmarks for histone modification during mitosis.

## 1 Introduction

During the cell division process, gene transcription must be initially terminated and then reactivated once cell division is complete. However, the specific mechanism and factors controlling this process of transcription regulation remain unclear. Since it would be highly time- and energy-consuming to mark all genes that need to be transcribed from scratch after each cycle of cell division, it has been proposed that genes that need to be transcribed are “bookmarked” to easily recover these positions for reactivation [Festuccia et al.(2017)Festuccia, Gonzalez, Owens, and Navarro, Bellec et al.(2018)Bellec, Radulescu, and Lagha, Zaidi et al.(2018)Zaidi, Nickerson, Imbalzano, Lian, Stein, and Stein, Teves et al.(2016)Teves, An, Hansen, Xie, Darzacq, and Tjian]. Despite several proposals, the actual mechanism and nature of these “bookmarks” have not yet been identified. [John and Workman(1998)] suggested that condensed mitotic chromosomes can act as bookmarks, some histone modifications were suggested to serve as these bookmarks [Wang and Higgins(2013), Kouskouti and Talianidis(2005), Chow et al.(2005)Chow, Georgiou, Szutorisz, Maia e Silva, Pombo, Barahona et al.], and some transcription factors (TFs) have also been identified as potential bookmarks [Dey et al.(2000)Dey, Ellenberg, Farina, Coleman, Maruyama, Sciortino et al., Kadauke et al.(2012)Kadauke, Udugama, Pawlicki, Achtman, Jain, Cheng et al., Xing et al.(2005)Xing, Wilkerson, Mayhew, Lubert, Skaggs, Goodson et al., Christova and Oelgeschläger(2001), Festuccia et al.(2016)Festuccia, Dubois, Vandormael-Pournin, Tejeda, Mouren, Bessonnard et al.].

Recently, [Kang et al.(2020)Kang, Shokhirev, Xu, Chandran, Dixon, and Hetzer] suggested that histone 3 methylation or trimethylation at lysine 4 (H3K4me1 and H3K4me3, respectively) can act as a “bookmark” to identify genes to be transcribed, and that a limited number of TFs might act as bookmarks. However, there has been no comprehensive search of candidate “bookmark” TFs based on large-scale datasets.

We here propose a novel computational approach to search for TFs that might act as “bookmarks” during mitosis, which involves tensor decomposition (TD)-based unsupervised feature extraction (FE) (Fig. 1). In brief, after fragmenting the whole genome into DNA regions of 25,000 nucleotide, the histone modifications within each region were summed. In this context, each DNA region is considered a tensor and various singular-value vectors associated with either the DNA region or experimental conditions (e.g., histone modification, cell line, and cell division phase) are derived. After investigating singular-value vectors attributed to various experimental conditions, the DNA regions with significant associations of singular-value vectors attributed to various experimental conditions were selected as potentially biologically relevant regions. The genes included in the selected DNA regions were then identified and uploaded to the enrichment server Enrichr to identify TFs that target the genes. To our knowledge, this is the first method utilizing a TD-based unsupervised FE approach in a fully unsupervised fashion to comprehensively search for possible candidate bookmark TFs.

**Fig 1.**
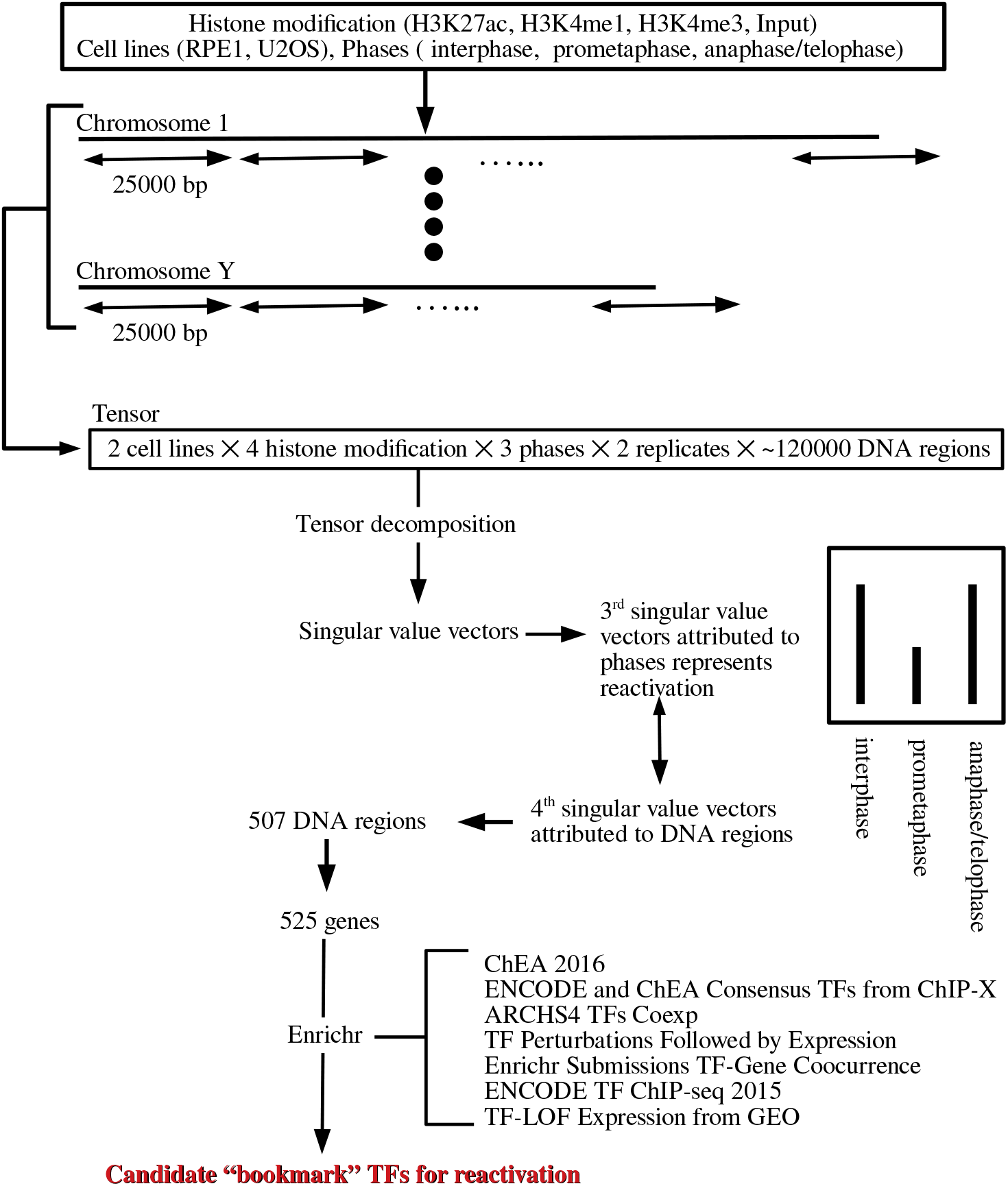
Flow chart of analyses performed in this study

## 2 Materials and Methods

### 2.1 Histone modification

The whole-genome histone modification profile was downloaded from the Gene Expression Omnibus (GEO) GSE141081 dataset. Sixty individual files (with extension.bw) were extracted from the raw GEO file. After excluding six CCCTC-binding factor (CTCF) chromatin immunoprecipitation-sequencing files and six 3rd replicates of histone modification files, a total of 48 histone modification profiles were retained for analysis. The DNA sequences of each chromosome were divided into 25,000-bp regions. Note that the last DNA region of each chromosome may be shorter since the total nucleotide length does not always divide into equal regions of 25,000. Histone modifications were then summed in each DNA region, which was used as the input value for the analysis. In total, *N* = 123, 817 DNA regions were available for analysis. Thus, with approximately 120, 000 regions of 25, 000 bp each, we covered the approximate human genome length of 3 × 10^9^.

### 2.2 Tensor Data Representation

Histone modification profiles were formatted as a tensor, *x*_*ijkms*_ ∈ R^*N ×*2*×*4*×*3*×*2^, which corresponds to the *k*th histone modification (*k* = 1: acetylation, H3K27ac; *k* = 2: H3K4me1; *k* = 3: H3K4me3; and *k* = 4 :Input) at the *i*th DNA region of the *j*th cell line (*j* = 1: RPE1 and *j* = 2: USO2) at the *m*th phase of the cell cycle(*m* = 1: interphase, *m* = 2: prometaphase, and *m* = 3: anaphase/telophase) of the *s*th replicate (*s* = 1, 2). *x*_*ijkms*_ was normalized as 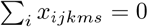 and 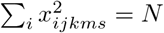 (Table 1). There are two biological replicates for each of the combinations of one of cell lines (either RPE1 or USO2), one of ChIP-seq (either acetylation or H3Kme1 or H3Kme4 or inout), and one of three cell cycle phases.

**Table 1.** Combinations of experimental conditions. Individual conditions are associated with two replicates

| Phases | Histone modifications |  |  |  |  |  |  |  |
| --- | --- | --- | --- | --- | --- | --- | --- | --- |
|  | Cell lines |  |  |  |  |  |  |  |
|  | H3K27ac |  | H3K4me1 |  | H3K4me3 |  | Input |  |
|  | RPE1 | U2OS | RPE1 | U2OS | RPE1 | U2OS | RPE1 | U2OS |
| interphase | ○ | ○ | ○ | ○ | ○ | ○ | ○ | ○ |
| prometaphase | ○ | ○ | ○ | ○ | ○ | ○ | ○ | ○ |
| anaphase/telophase | ○ | ○ | ○ | ○ | ○ | ○ | ○ | ○ |

### 2.3 Tensor Decomposition

Higher-order singular value decomposition (HOSVD) [Taguchi(2020)] was applied to *x*_*ijkms*_ to obtain the decomposition

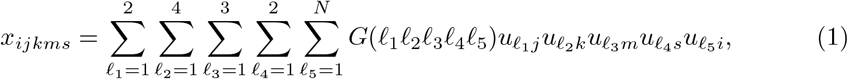

where *G* ∈ R^2*×*4*×*3*×*2*×N*^ is the core tensor, and 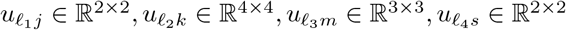 and 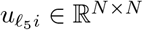 are singular-value vector matrices, which are all orthogonal matrices. The reason for using the complete representation instead of the truncated representation of TD is that we employed HOSVD to compute TD. In HOSVD, the truncated representation is equal to that of the complete representation; i.e., 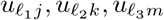 and 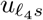 are not altered between the truncated and the full representation. For more details, see [Taguchi(2020)].

Here is a summary on how to compute eq. (1) using the HOSVD algorithm, although it has been described in detail previously [Taguchi(2020)]. At first, *x*_*ijkms*_ is unfolded to a matrix, *x*_*i*(*jkms*)_ ∈ R^*N ×*48^. Then SVD is applied to get

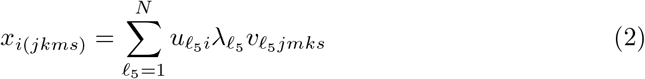

Then, only 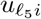 is retained, and 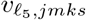 is discarded. Similar procedures are applied to *x*_*ijkms*_ by replacing *i* with one of *j, k, m, s* in order to get 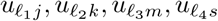. Finally, *G* can be computed as

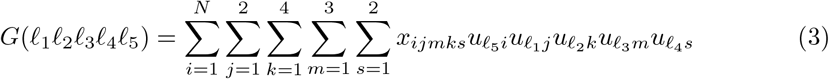

### 2.4 TD-based unsupervised FE

Although the method was fully described in a recently published book [Taguchi(2020)], we summarize the process of selecting genes starting from the TD.

- To identify which singular value vectors attributed to samples (e.g., cell lines, type of histone modification, cell cycle phase, and replicates) are associated with the desired properties (e.g., “not dependent upon replicates or cell lines,” “represents re-activation,” and “distinct between input and histone modifications”), the number of singular value vectors selected are not decided in advance, since there is no way to know how singular value vectors behave in advance, because of the unsupervised nature of TD.
- To identify which singular value vectors attributed to genomic regions are associated with the desired properties described above, core tensor, *G*, is investigated. We select singular value vectors attributed to genomic regions that share *G* with larger absolute values with the singular value vectors selected in the process mentioned earlier, because these singular value vectors attributed to genomic regions are likely associated with the desired properties.
- Using the selected singular value vectors attributed to genomic regions, those associated with the components of singular value vectors with larger absolute values are selected, because such genomic regions are likely associated with the desired properties. Usually, singular value vectors attributed to genomic regions are assumed to obey Gaussian distribution (null hypothesis), and *P* -values are attributed to individual genomic regions. *P* -values are corrected using multiple comparison correction, and the genomic regions associated with adjusted *P* -values less than the threshold value are selected.
- There are no definite ways to select singular value vectors. The evaluation can only be done using the selected genes. If the selected genes are not reasonable, alternative selection of singular value vectors should be attempted. When we cannot get any reasonable genes, we abort the procedure.

To select the DNA regions of interest (i.e., those associated with transcription reactivation), we first needed to specify the singular-value vectors that are attributed to the cell line, histone modification, phases of the cell cycle, and replicates with respect to the biological feature of interest, transcription reactivation. Consider selection of a specific index set 𝓁_1_, 𝓁_2_, 𝓁_3_, 𝓁_4_ as one that is associated with biological features of interest, we then select 𝓁_5_ that is associated with *G* with larger absolute values, since singular-value vectors 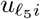 with 𝓁_5_ represent the degree of association between individual DNA regions and reactivation. Using 𝓁_5_, we attribute *P* -values to the *i*th DNA region assuming that 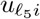 obeys a Gaussian distribution (null hypothesis) using the *χ*^2^ distribution

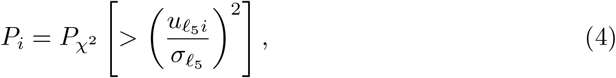

where 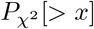 is the cumulative *χ*^2^ distribution in which the argument is larger than *x*, and 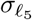 is the standard deviation. *P* -values are then corrected by the BH criterion [Taguchi(2020)], and the *i*th DNA region associated with adjusted *P* -values less than 0.01 were selected as those significantly associated with transcription reactivation.

Algorithm displayed with mathematical formulas can be available in Fig. 2.

**Fig 2.**
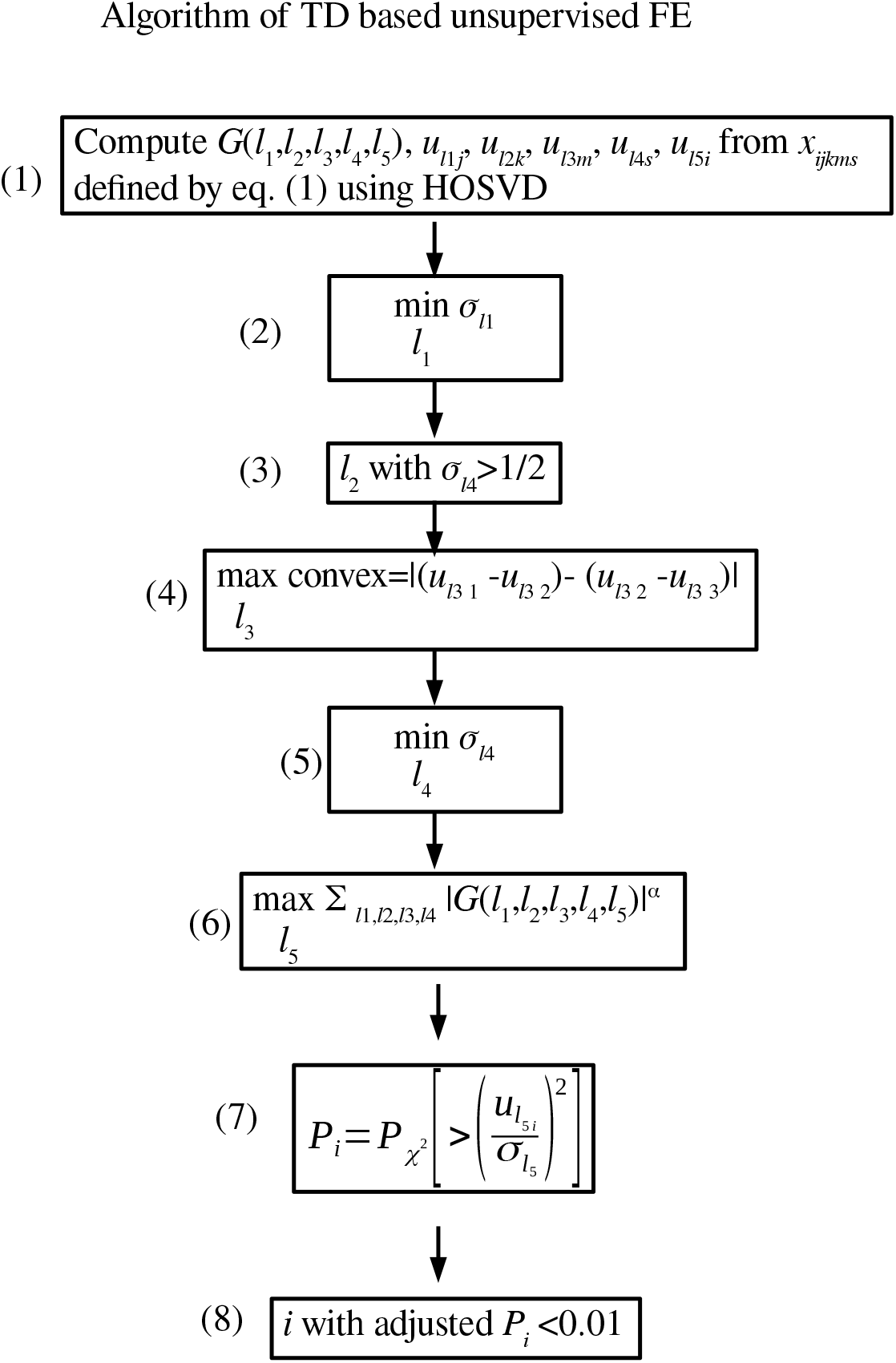
Algorithm of TD based unsupervised FE. (1) Perform TD to derive *G*(𝓁_1_, 𝓁_2_, 𝓁_3_, 𝓁_4_, 𝓁_5_). (2) Select 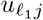 that takes constant values between two cell lines as much as possible. (3) Select 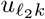 that has distinct values for Histone modification toward inputs. (4) Select 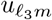 that represents reactivation during three cell cycle phases as much as possible. (5) Select 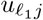 that takes constant values between two biological replicates as much as possible. (6) Select 𝓁 _5_ associated with *G* having largest absolute values given 𝓁_1_, 𝓁_2_, 𝓁_3_, 𝓁_4_ (7) Attribute *P* -values to *i*s with assuming that 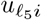 obeys Gaussian distribution (Null hypothesis). (8) Select *i*s associated with adjusted *P* -values less than 0.01.

### 2.5 Enrichment analysis

Gene symbols included in the selected DNA regions were retrieved using the biomaRt package [Durinck et al.(2009)Durinck, Spellman, Birney, and Huber] of R [R Core Team(2019)] based on the hg19 reference genome. The selected gene symbols were then uploaded to Enrichr [Kuleshov et al.(2016)Kuleshov, Jones, Rouillard, Fernandez, Duan, Wang et al.] for functional annotation to identify their targeting TFs.

### 2.6 DESeq2

When DESeq2 [Love et al.(2014)Love, Huber, and Anders] was applied to the present data set, six samples within each cell lines measured for three cell cycles and associated with two replicates were considered. Three cell cycles were regarded to be categorical classes associated with no rank order since we would like to detect not monotonic change between cell cycles but re-activation during them. All other parameters are defaults. Counts less than 1.0 were truncated so as to have integer values (e.g., 1400.53 was converted to 1400).

### 2.7 Csaw

Since csaw [Lun and Smyth(2015)] required bam files not available in GEO, we first mapped 60 fastq files to hg38 human genome using bowtie2 [Langmead and Salzberg(2012)] where 60 fastq files in GEO ID GSE141081 were downloaded from SRA. Sam files generated by bowtie2 were converted and indexed by samtools [Li et al.(2009)Li, Handsaker, Wysoker, Fennell, Ruan, Homer et al.] and sorted bam files were generated. Generated bam files that correspond to individual combinations of cell lines and ChIP-seq were loaded into csaw in order to identify differential binding among three cell cycle phases.

### 2.8 Identification of overlapping regions between peak call

We retrieved 36 peak call data set (with extension peaks.txt.gz) that correspond to 48 Chip-Seq files with excluding 12 input files. Starting from these 48 peak call files, using `findOverlapsOfPeaks` function included in ChIPpeakAnno package in R, we selected overlap regions step by step as follows

- Identify overlap regions between two biological replicates; this results in 9 regions for U2OS cell lines and RPE1 cell lines, respectively, in total 18 peak calls.
- Identify overlap regions among three cell cycles; retrieve regions commonly expressed in three cell cycle phases for H3K4me1 and H3K4me3 whereas those expressed only in interphase and anaphase/telophase; this results in three regions, each of which was attributed to H3K4me1, H3K27ac, or H3K4me3, for U2OS cell lines and RPE1 cell lines, respectively, in total 6 peak calls.
- Identify overlap between 6 peak calls.

This process was illustrated in Fig. 3.

**Fig 3.**
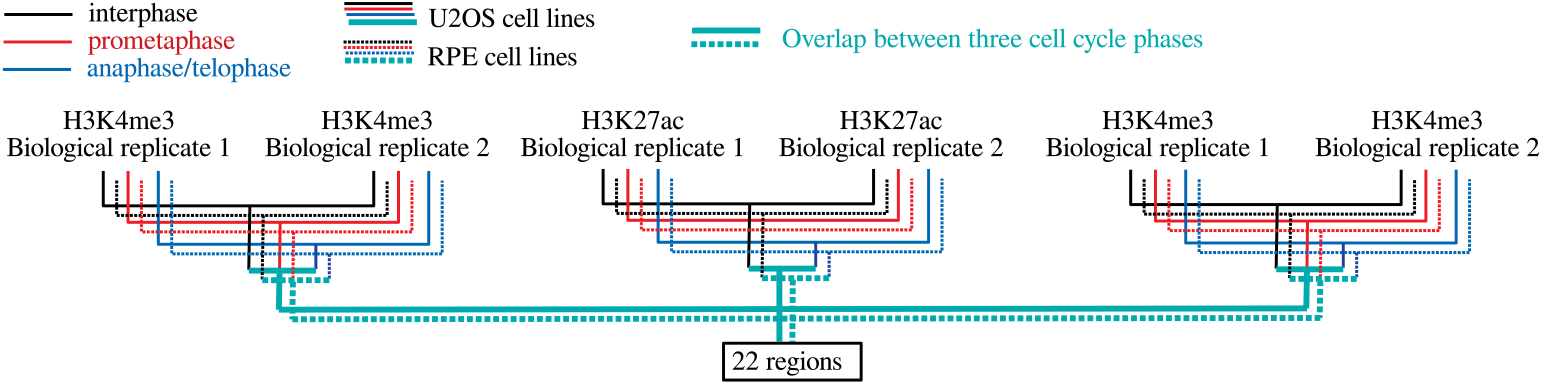
Flow chart on how we integrated peak call files.

## 3 Results and Discussion

We first attempted to identify which singular-value vector is most strongly attributed to transcription reactivation among the vectors for cell line 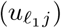 histone modification 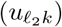 cell cycle phase 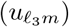 and replicate 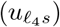 (Fig. 4). First, we considered phase dependency. Fig. 5 shows the singular-value vectors 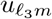 attributed to cell cycle phases. In the case that there are a set of genes that share some dependence, singular value vectors reflect their mean behaviour. Specifically, singular value vectors act as some kind of pseudo representative genes. Thus, by investigating singular value vectors, we can find what kind of cell cycle dependence can appear in the group of genes. Since the reactivation means that being expressive in inter and ana/telophases whereas not expressive in prometapahse, singular value vectors supposed to be related to be reactivation take opposite signs between inter/ana/telophased and prometaphase. Thus, *u*_3*m*_ are most likely associated with reactivation. Although *u*_2*m*_ and *u*_3*m*_ were associated with reactivation, we further considered only *u*_3*m*_ since it showed a more pronounced reactivation profile. Next, we investigated singular-value vectors 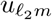 attributed to histone modification (Fig. 6). There was no clearly interpretable dependence on histone modification other than for *u*_1*k*_ which represents the lack of histone modification, since the values for H3K27ac, H3K4me1, and H3K4me3 were equivalent to the Input value that corresponds to the control condition; thus, *u*_2*k*_, *u*_3*k*_, and *u*_4*k*_ were considered to have equal contributions for subsequent analyses. By contrast, since *u*_1*j*_ and *u*_1*s*_ showed no dependence on cell line and replicates, respectively, we selected these vectors for further downstream analyses (Fig. 7).

**Fig 4.**
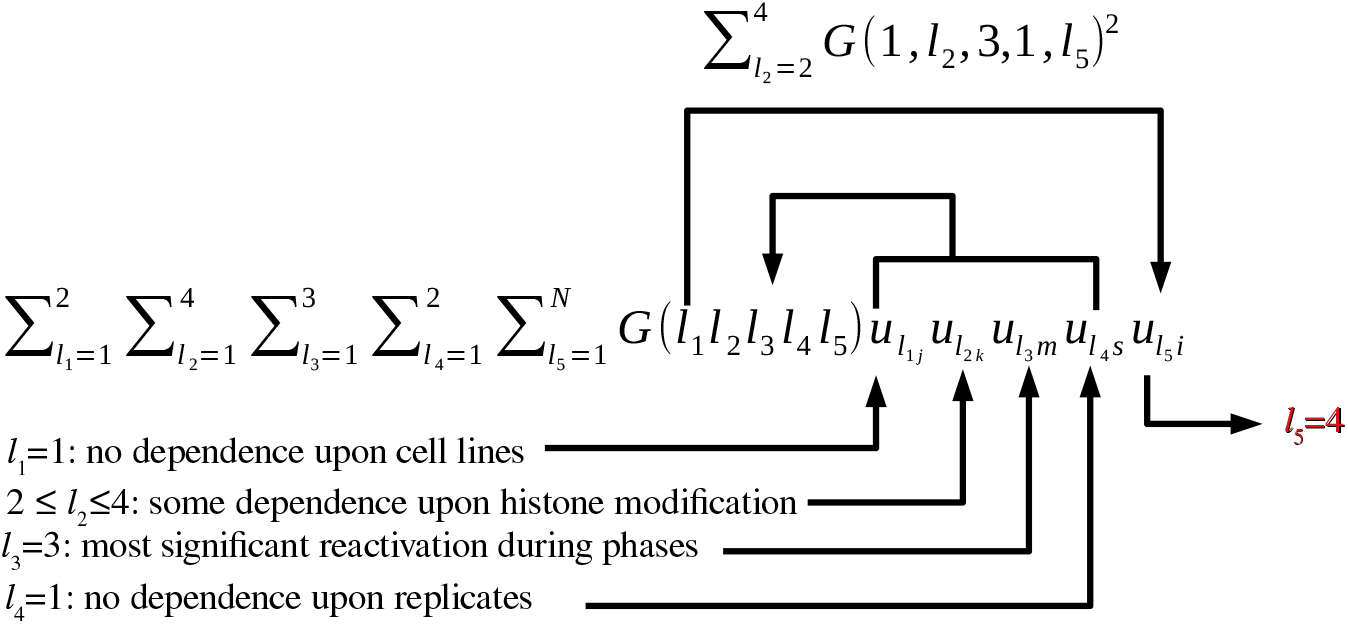
Schematic of the process for selecting *u*_4*i*_ to be used for DNA region selection.

**Fig 5.**
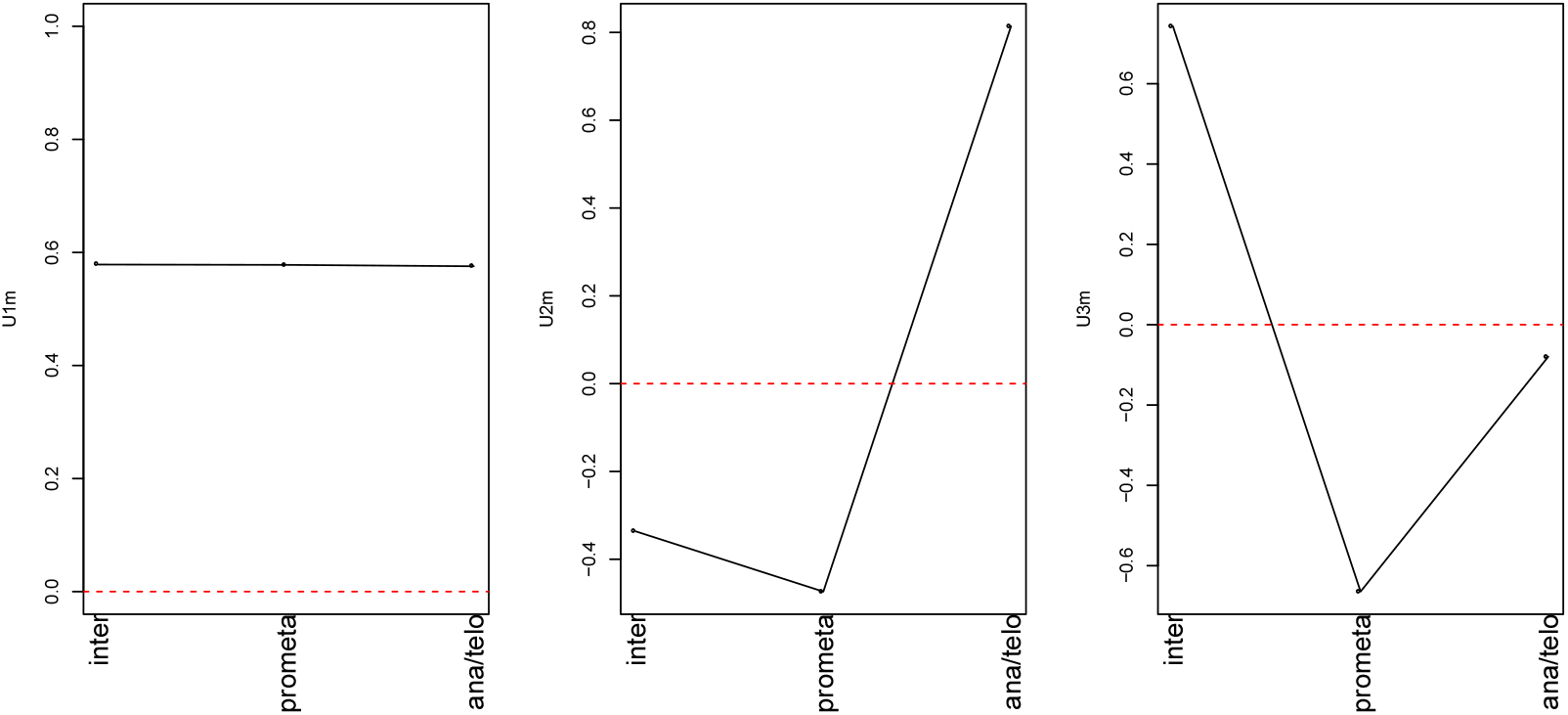
Singular-value vectors associated with cell cycle phase. Left: *u*_1*m*_, middle: *u*_2*m*_, right: *u*_3*m*_

**Fig 6.**
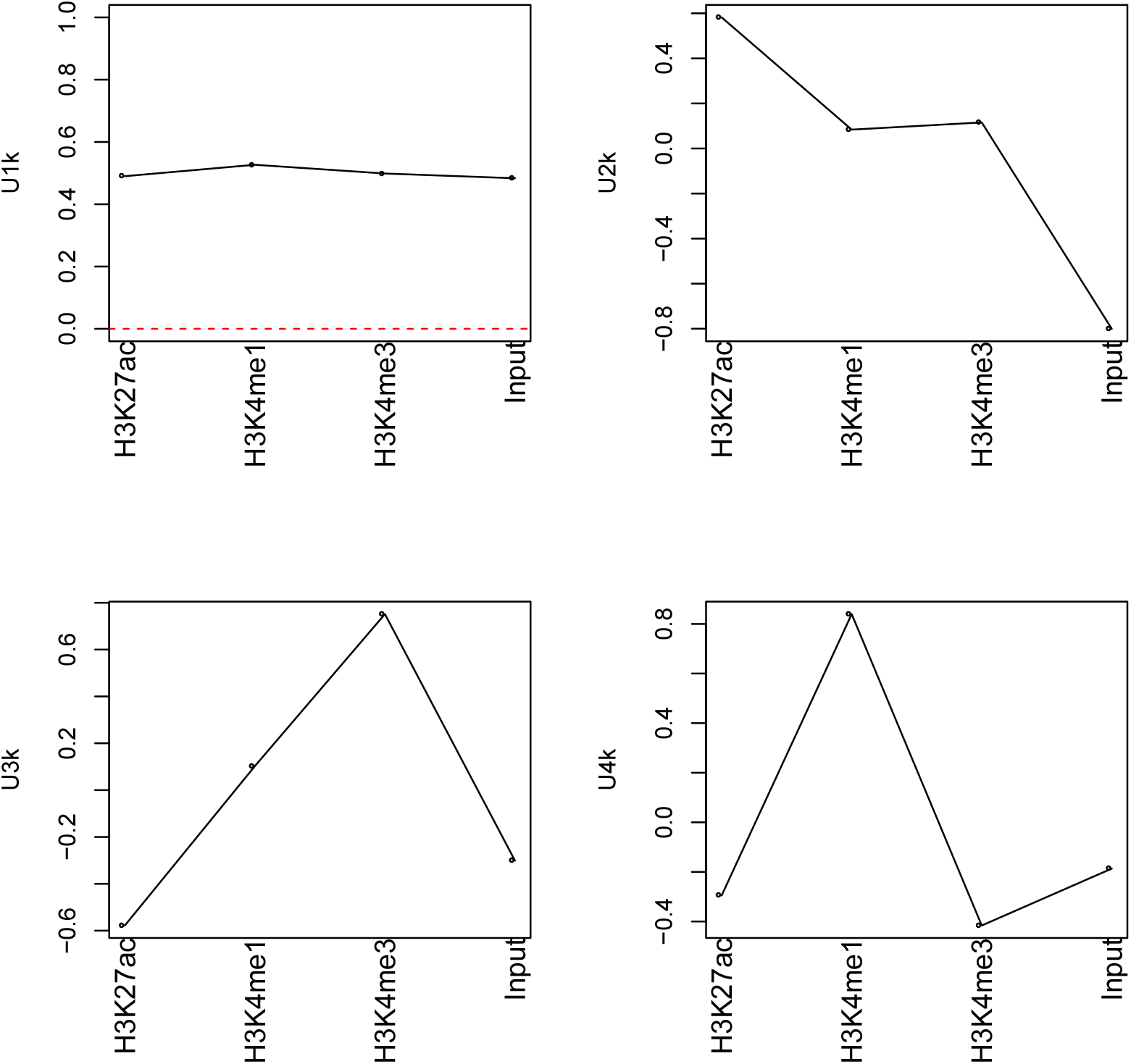
Singular-value vectors associated with histone modification. Upper left: *u*_1*k*_, upper right: *u*_2*k*_, lower left: *u*_3*k*_, lower right: *u*_4*k*_

**Fig 7.**
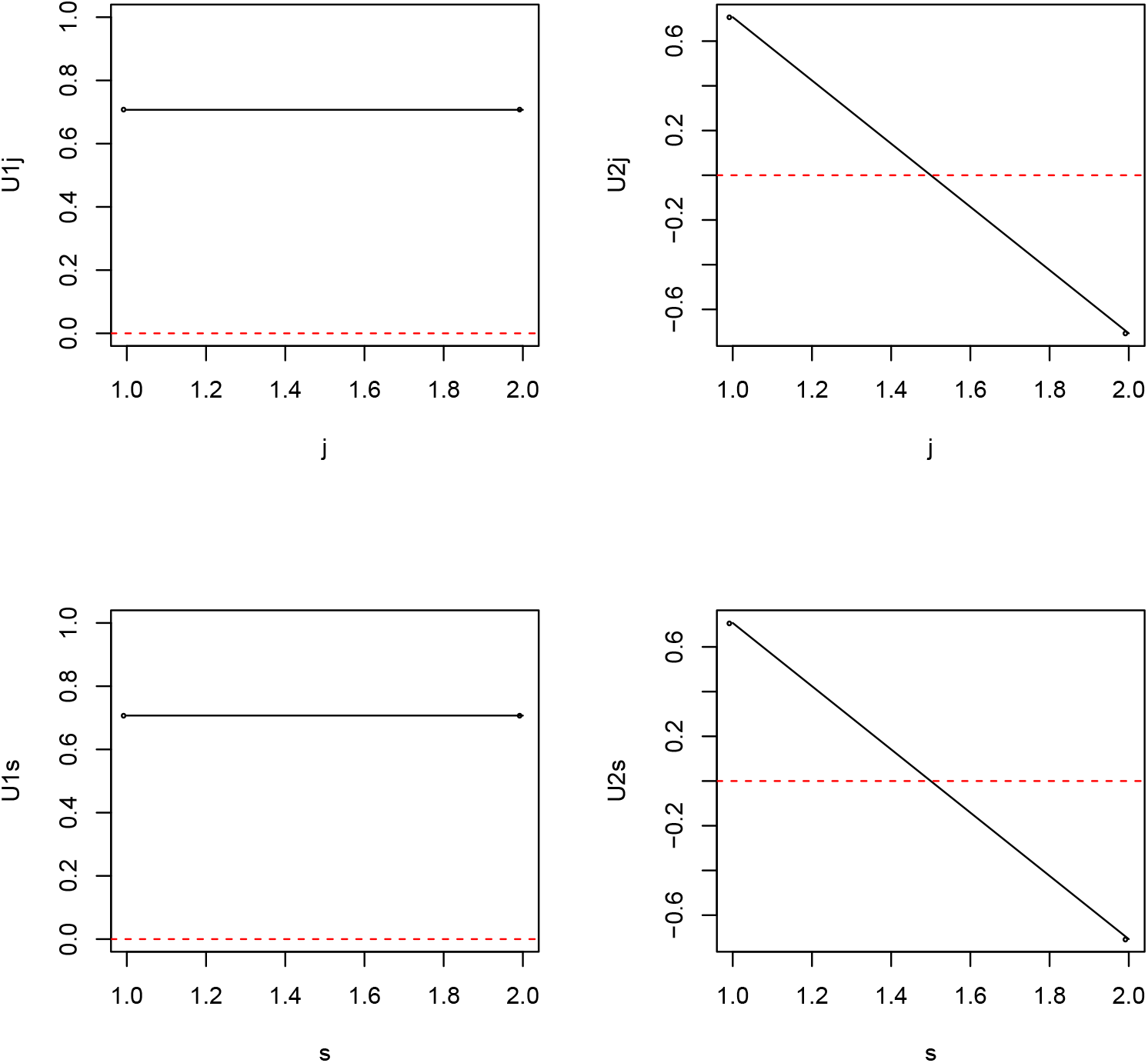
Dependence of vectors on cell line (j) and replicate (s). Top left: *u*_1*j*_, top right: *u*_2*j*_, bottom left: *u*_1*s*_, bottom right: *u*_2*s*_

Finally, we evaluated which vector 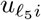 had a larger 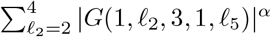 *α* = 1, 2, 3 (Fig. 8); in this case, we calculated the squared sum for 2 ≤ 𝓁_2_ ≤ 4 to consider them equally. Although we do not have any definite criterion to decide *α* uniquely, since 𝓁_5_ = 4 always takes largest values for *α* ≥ 1, 𝓁_5_ = 4 was further employed. The *P* -values attributed to the *i*th DNA regions were calculated using eq. (4), resulting in selection of 507 DNA regions associated with adjusted *P* -values less than 0.01.

**Fig 8.**
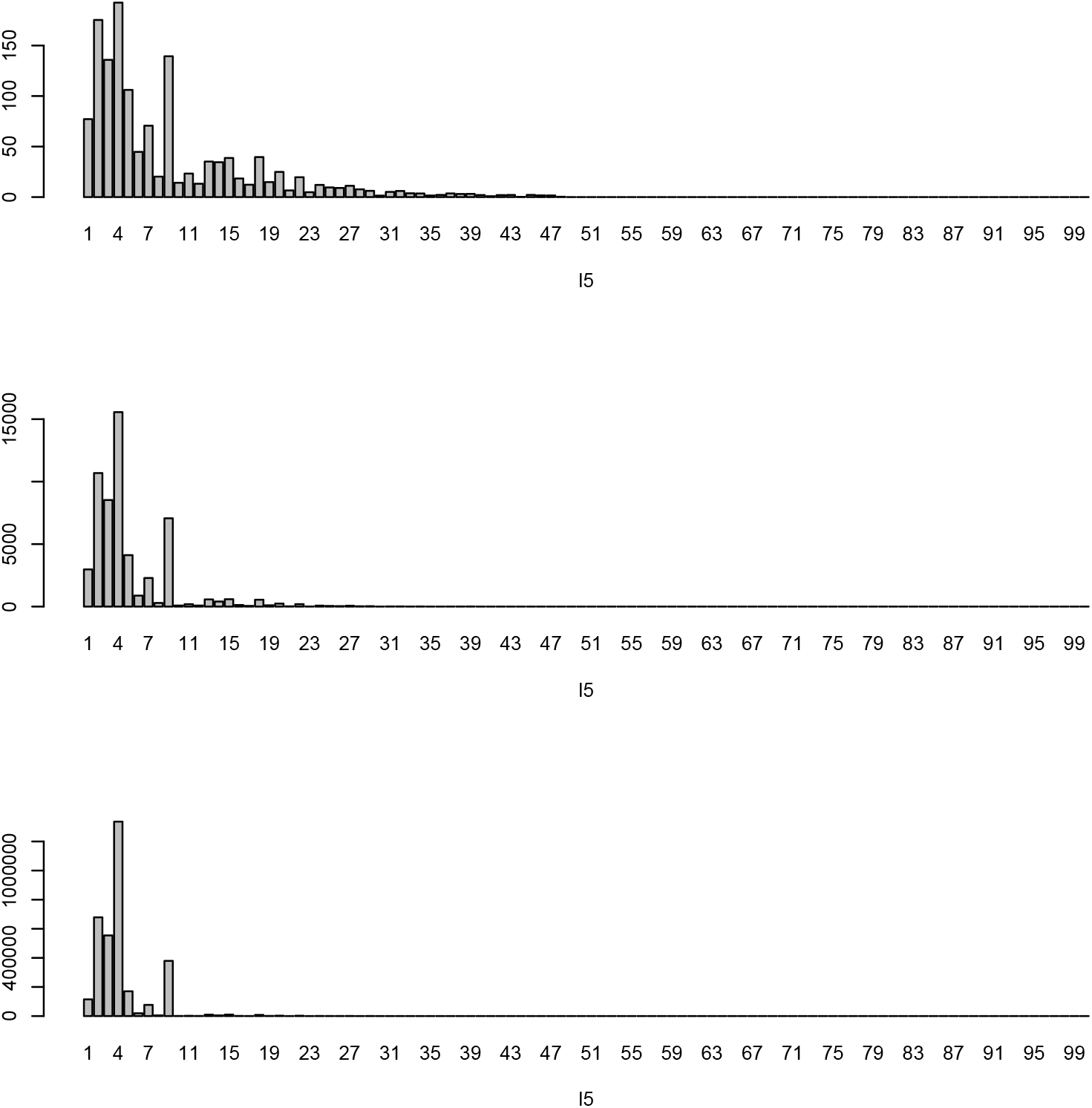
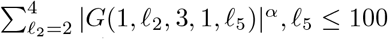. Because of HOSVD algorithm, *G*(𝓁_1_, 𝓁_2_, 𝓁_3_, 𝓁_4_, 𝓁_5_) = 0 for 𝓁_5_ *>* 2 × 4 × 3 × 2 = 48. *α* = 1 (Top), 2 (middle), and 3 (bottom).

We next checked whether histone modification in the selected DNA regions was associated with the following transcription reactivation properties:

1. H3K27ac should have larger values in interphase and anaphase/telophase than in prometaphase, as the definition of reactivation.
2. H3K4me1 and H3K4me3 should have constant values during all phases of the cell cycle, as the definition of a “bookmark” histone modification
3. H3K4me1 and H3K4me3 should have larger values than the Input; otherwise, they cannot be regarded to act as “bookmarks” since these histones must be significantly modified throughout these phases.

To check whether the above criteria are fulfilled, we applied six *t* tests to histone modifications in the 507 selected DNA regions (Table 2). The results clearly showed that histone modifications in the 507 selected DNA regions satisfied the requirements for transcription reactivation; thus, our strategy could successfully select DNA regions that demonstrate reactivation/bookmark functions of histone modification.

**Table 2.**
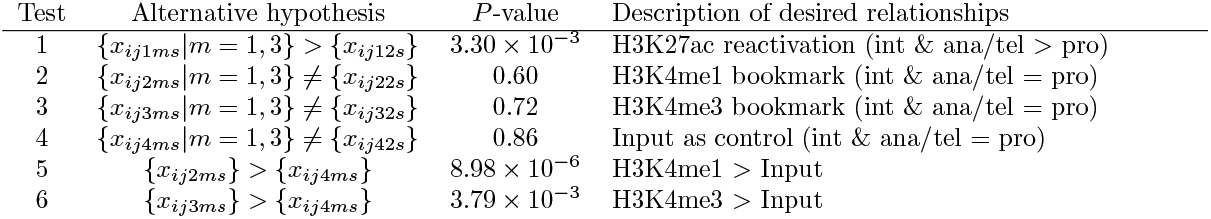
Hypotheses for *t* tests applied to histone modification in the selected 507 DNA regions. The null hypothesis was that the inequality relationship of the alternative hypothesis is replaced with an equality relationship. int: interphase, ana: anaphase, tel: telophase, pro: prometaphase.

After confirming that selected DNA regions are associated with targeted reactivation/bookmark features, we queried all gene symbols contained within these 507 regions to the Enrichr server to identify TFs that significantly target these genes. These TFs were considered candidate bookmarks that remain bound to these DNA regions throughout the cell cycle and trigger reactivation in anaphase/telophase (i.e., after cell division is complete). Table 3 lists the TFs associated with the selected regions at adjusted *P* -values less than 0.05 in each of the seven categories of Enrichr.

**Table 3.** Number of transcription factors (TFs) associated with adjusted *P* -values less than 0.05 in various TF-related Enrichr categories

| | | Adjusted $P$-values | |
| --- | --- | --- | --- |
| Terms |  | > 0.05 | < 0.05 |
| (I) | ChEA 2016 | 537 | 97 |
| (II) | ENCODE and ChEA Consensus TFs from ChIP-X | 91 | 12 |
| (III) | ARCHS4 TFs Coexp | 1533 | 54 |
| (IV) | TF Perturbations Followed by Expression | 1577 | 346 |
| (V) | Enrichr Submissions TF-Gene Cooccurrence | 587 | 1135 |
| (VI) | ENCODE TF ChIP-seq 2015 | 788 | 28 |
| (VII) | TF-LOF Expression from GEO | 239 | 11 |

Among the many TFs that emerged to be significantly likely to target genes included in the 507 DNA regions selected by TD-based unsupervised FE, we here focus on the biological functions of TFs that were also detected in the original study suggesting that TFs might function as histone modification bookmarks for transcription reactivation [Kang et al.(2020)Kang, Shokhirev, Xu, Chandran, Dixon, and Hetzer]. RUNX was identified as an essential TF for osteogenic cell fate, and has been associated with mitotic chromosomes in multiple cell lines, including Saos-2 osteosarcoma cells and HeLa cells (Young et al. 2007). Table 4 shows the detection of RUNX family TFs in seven TF-related categories of Enrichr; three RUNX TFs were detected in at least one of the seven TF-related categories. In addition, TEADs (Kegelman et al. 2018), JUNs [Wagner(2002)], FOXOs [Rached et al.(2010)Rached, Kode, Xu, Yoshikawa, Paik, DePinho et al.], and FosLs citepKang01072020 were reported to regulate osteoblast differentiation. Tables 5, 6,7, and 8 show that two TEAD TFs, three JUN TFs, four FOXO TFs, and two FOSL TFs were detected in at least one of the seven TF-related categories in Enrichr, respectively.

**Table 4.**
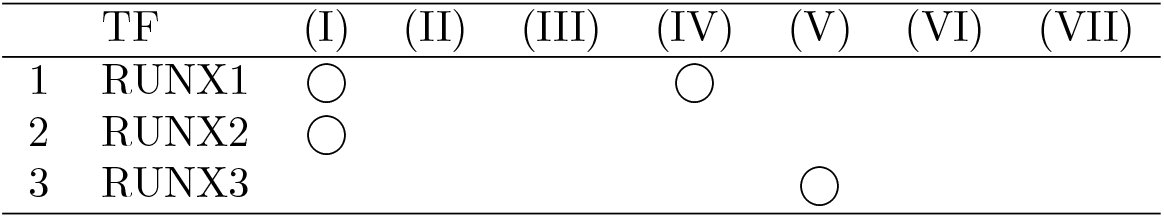
Identification of RUNX transcription factor (TF) family members within seven TF-related categories in Enrichr. Roman numerals correspond to the first column in Table 3.

| TF | (I) | (II) | (III) | (IV) | (V) | (VI) | (VII) |
| --- | --- | --- | --- | --- | --- | --- | --- |
| 1 RUNX1 | ○ |  |  | ○ |  |  |  |
| 2 RUNX2 | ○ |  |  |  |  |  |  |
| 3 RUNX3 |  |  |  |  | ○ |  |  |

**Table 5.** Identification of TEAD transcription factor (TF) family members within seven TF-related categories in Enrichr. Roman numerals correspond to the first column in Table 3.

| TF | (I) | (II) | (III) | (IV) | (V) | (VI) | (VII) |
| --- | --- | --- | --- | --- | --- | --- | --- |
| 1 TEAD4 | ○ |  |  |  |  | ○ |  |
| 2 TEAD3 |  |  | ○ |  |  |  |  |

**Table 6.** Identification of JUN transcription factor (TF) family members within seven TF-related categories in Enrichr. Roman numerals correspond to the first column in Table 3.

| TF | (I) | (II) | (III) | (IV) | (V) | (VI) | (VII) |
| --- | --- | --- | --- | --- | --- | --- | --- |
| 1 JUN | ○ |  |  | ○ | ○ | ○ |  |
| 2 JUND | ○ |  |  | ○ | ○ | ○ |  |
| 3 JUNB |  |  |  | ○ | ○ |  |  |

**Table 7.**
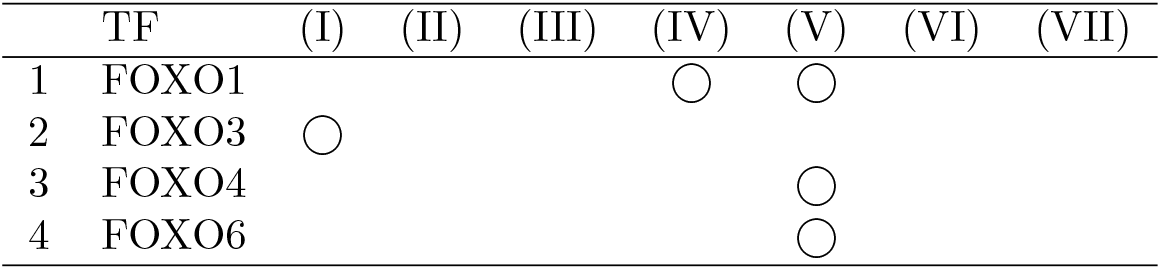
Identification of FOXO transcription factor (TF) family members within seven TF-related categories in Enrichr. Roman numerals correspond to the first column in Table 3.

**Table 8.**
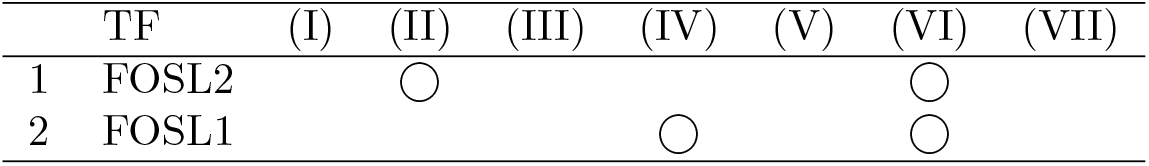
Identification of FosL transcription factor (TF) family members within seven TF-related categories in Enrichr. Roman numerals correspond to the first column in Table 3.

Other than these five TF families reported in the original study [Kang et al.(2020)Kang, Shokhirev, Xu, Chandran, Dixon, and Hetzer], the TFs detected most frequently within seven TF-related categories in Enrichr were as follows (Table 9): GATA2 [Kala et al.(2009)Kala, Haugas, Lilleväli, Guimera, Wurst, Salminen et al.], ESR1 [Kato and Ogawa(1994)], TCF21 [Kim et al.(2017)Kim, Pjanic, Nguyen, Miller, Iyer, Liu et al.], TP53 [Ha et al.(2007)Ha, Baek, Kim, Jeong, Kim, McKeon et al.], WT1 [Shandilya and Roberts(2015)], NFE2L2 (also known as NRF2 [Martin-Hurtado et al.(2019)Martin-Hurtado, Martin-Morales, Robledinos-Antón, Blanco, Palacios-Blanco, Lastres-Becker et al.]), GATA1 [Kadauke et al.(2012)Kadauke, Udugama, Pawlicki, Achtman, Jain, Cheng et al.], and GATA3 [Shafer et al.(2017)Shafer, Nguyen, Tremblay, Viala, Béland, Bertos et al.]. All of these TFs have been reported to be related to mitosis directly or indirectly, in addition to JUN and JUND, which are listed in Table 6. This further suggests the suitability of our search strategy to identify transcription reactivation bookmarks.

**Table 9.** Top 10 most frequently listed transcription factor (TF) families (at least four, considered the majority) within seven TF-related categories in Enrichr. Roman numerals correspond to the first column in Table 3.

|  | TF | (I) | (II) | (III) | (IV) | (V) | (VI) | (VII) |
| --- | --- | --- | --- | --- | --- | --- | --- | --- |
| 1 | GATA2 | ○ | ○ |  | ○ | ○ | ○ |  |
| 2 | ESR1 | ○ | ○ |  | ○ | ○ | ○ |  |
| 3 | TCF21 | ○ |  | ○ | ○ | ○ |  |  |
| 4 | TP53 | ○ | ○ |  | ○ | ○ |  |  |
| 5 | JUN | ○ |  |  | ○ | ○ | ○ |  |
| 6 | JUND | ○ |  |  | ○ | ○ | ○ |  |
| 7 | WT1 | ○ |  |  | ○ | ○ |  | ○ |
| 8 | NFE2L2 | ○ | ○ |  | ○ | ○ |  |  |
| 9 | GATA1 | ○ | ○ |  | ○ | ○ |  |  |
| 10 | GATA3 |  |  |  | ○ | ○ | ○ | ○ |

One might wonder why we did not compare our methods with the other methods. As can be seen in Table 1, there are only two samples each in as many as 24 categories. Therefore, it is difficult to apply standard statistical tests for pairwise comparisons between two groups including only two samples. In addition, the number of features, *N*, which is the number of genomic regions in this study, is as many as 1,23,817, which drastically reduces the significance of each test if we consider multiple comparison criteria that increase *P* -values that reject the null hypothesis. Finally, only a limited number of pairwise comparisons are meaningful; for example, we are not willing to compare the amount of H3K4me1 in the RPE1 cell line at interphase with that of H3K27ac in the U2OS cell line at prometaphase. Therefore, usual procedures that deal with pairwise comparisons comprehensively, such as Tukey’s test, cannot be applied to the present data set as it is. In conclusion, we could not find any suitable method applicable to the present data set that has a small number of samples within each of as many as 24 categories, whereas the number of features is as many as 1,23,817.

In order to demonstrate inferiority of other method compared with our method, we applied DESeq2 [Love et al.(2014)Love, Huber, and Anders] to the present data set, although DESeq2 was designed to not ChIP-seq but RNA-seq. The outcome is disappointing as expected (Table 10) if it is compared with Table 2. First of all, there are no coincidences between two cell lines. Although there are as many as 4227 regions within which H3K4me1 is distinct among three cell cycle phases when RPE1 is considered, there were no regions associated with distinct H3K4me1 when U2OS was considered. In addition to this, although only H3K27ac among three histone modifications measured is expected to be distinct during three cell cycle phases, other histone modifications are sometimes detected as distinct during three cell cycle phases. Finally, the number of genomic regions considered in each comparison varies, since DESeq2 automatically discarded regions associated with low variance among distinct classes. The reason why there are no regions associated with distinct histone modification for Input and H3K4me1 when RPE1 was considered is definitely because almost all genomic regions were considered for these two comparisons; too many comparisons increase the *P* -values because of multiple comparison corrections. On the other hand, our proposed TD based unsupervised FE can deal with all of the genomic regions, which resulted in more stable outcomes. Thus, it is obvious that DESeq2 was inferior to TD based unsupervised FE when it is applied to the present data set.

**Table 10.** The performances achieved by DESeq2 applied to the present data set. Adjp: adjusted *P* -values computed by DESeq2

|  | RPE1 |  | U2OS |  |
| --- | --- | --- | --- | --- |
|  | Adjp > 0.01 | Adjp < 0.01 | Adjp > 0.01 | Adjp < 0.01 |
| H3K27ac | 30649 | 1829 | 28849 | 1425 |
| H3K4me1 | 113784 | 0 | 52323 | 4227 |
| H3K4me3 | 26420 | 8259 | 24359 | 1559 |
| Input | 112976 | 0 | 5995 | 196 |

One might still wonder if it is because of usage of DESeq2 not designed specific to ChIP-seq data. In order to confirm this point, we sought integrated approaches designed specific to treatment of ChIP-seq data. In addition, we need some approaches that enable us not only pairwise comparison but also comparisons among more than two categories, since we have to compare among three cell cycle phases, i.e., terphase, prometaphase, and anaphase/telophase. There are not so many approaches satisfying these conditions [Wu et al.(2015)Wu, Bittencourt, Stallcup, and Siegmund, Steinhauser et al.(2016)Steinhauser, Kurzawa, Eils, and Herrmann, Tu and Shao(2017)]. For example, although DBChIP [Liang and Keleş(2011)] was designed to treat ChIP-seq data set, since it was designed to be specific to TF binding, it required to input single nucleotide positions where binding proteins bind, Thus, it is not applicable to histone modification measurements where not binding points but binding regions are provided. On the other hand, although DiffBind [Stark and Brown(2011)] was designed to deal with histone modification, it can accept only pairwise comparisions. SCIFER [Xu et al.(2014)Xu, Grullon, Ge, and Peng] can identify enrichment within single measurement compared with input experiment, MACS2 which is modified version of MACS [Zhang et al.(2008)Zhang, Liu, Meyer, Eeckhoute, Johnson, Bernstein et al.], can also accept only pairwise comaprisons, ODIN [Allhoff et al.(2014)Allhoff, Seré, Chauvistré, Lin, Zenke, and Costa] also can accept only pairwise comparisons, RSEG [Song and Smith(2011)] also can accept only pairwise comparisons, MAnorm [Shao et al.(2012)Shao, Zhang, Yuan, Orkin, and Waxman] also can accept only pairwise comparisons, HOMER [Heinz et al.(2010)Heinz, Benner, Spann, Bertolino, Lin, Laslo et al.] also can accept only pairwise comparisons, QChIPat [Liu et al.(2013)Liu, Yi, SV, Lan, Ma, Huang et al.] also can accept only pairwise comparisons, diffReps [Shen et al.(2013)Shen, Shao, Liu, Maze, Feng, and Nestler] also can accept only pairwise comparisons, MMDiff [Schweikert et al.(2013)Schweikert, Cseke, Clouaire, Bird, and Sanguinetti] also can accept only pairwise comparisons, PePr [Zhang et al.(2014)Zhang, Lin, Johnson, Rozek, and Sartor] does not perform even pairwise comparison. ChIPComp [Chen et al.(2015)Chen, Wang, Qin, and Wu] was tested toward only pairwise comparisons when it was applied to real data set. Although MultiGPS [Mahony et al.(2014)Mahony, Edwards, Mazzoni, Sherwood, Kakumanu, Morrison et al.] can deal with multiple files, they must be composed of condition A and its corresponding input vs condition B and its corresponding input, it cannot be applied to the present case composed of three cell cycle phases and their corresponding inputs. Thus as far as we investigated there are no approaches designed to be applicable to three independent conditions, each of which is composed of a pair of treated and input experiments.

This difficulty is because of two kinds of distinct differential binding analyses required (Fig. 9), one of which is the comparison between treated and input experiments and another of which is the comparison between two experimental conditions (e.g., patients versus healthy control, two different tissues) whereas they are easily performed in tensor representation as shown in the above. Nevertheless, in order to emphasize the inferiority of ChIP-seq specific pipeline aiming differential binding analysis toward TD based unsupervised FE, we considered csaw [Lun and Smyth(2015)] as a representative since it accepts, at least, not pairwise but comparisons among multiple conditions as performed by DESeq2 (Table 10). Table 11 shows the results. It is very disappointing as expected. For example, although H3K27ac is expected to support reactivation, differential binding region among distinct cell cycle phases in U2OS cell line is almost none (only 0.1 % of whole tested regions). Although H3K4me3 should not distinctly bind to chromosome among thee cell cycles since it is expected to play a role of bookmark, it distinctly binds to chromosomes among three cell cycle phases for two cell lines. These behaviours are very contrast to those in Table 2 which exhibits the expected differential/undifferential binding to chromosome. Thus, in conclusion, even if we employ pipelines specifically designed to ChIP-Seq data analyses, they cannot outperform the results obtained by TD based unsupervised FE.

**Table 11.** The performances achieved by csaw applied to the present data set. Adjp: adjusted *P* -values computed by csaw

|  | RPE1 |  | U2OS |  |
| --- | --- | --- | --- | --- |
|  | Adjp > 0.01 | Adjp < 0.01 | Adjp > 0.01 | Adjp < 0.01 |
| H3K27ac | 4127704 | 113803 | 4477318 | 6126 |
| H3K4me1 | 5552148 | 0 | 6060553 | 5 |
| H3K4me3 | 3054309 | 140962 | 2197717 | 27570 |
| Input | 3310106 | 0 | 5040796 | 0 |

**Fig 9.**
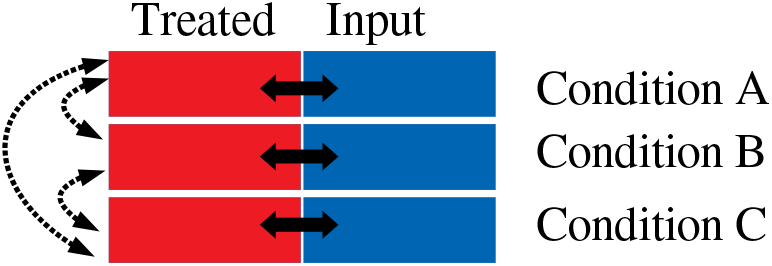
Schematics that illustrates the difficulty of differential binding analysis. In contrast to differential expression analysis that requires only inter conditions comparisons (displayed by broken bidirectional arrows), differential binding analysis requires additional intra conditions comparisons between treated and input experiment (displayed by bidirectional solid arrows). There are no pipelines that aim to identify differential binding considering simultaneously more than two conditions.

Since one might wonder why we have specifically used region length of 25,000 nucleic acid length, we discuss about it as follows.

- We have successfully used the region length [49, 50]. When started to employ this procedure, we tried multiple values and identified that it is most successful.
- Optimizing region length from studies to studies is not a good way to identify something biological. Region length should not be optimization parameters. If the optimal region length vary from studies to studies, we need to rationalize it. Nevertheless, the fact that employment region of 25,000 nucleic acid length was successful in three independent studies (including the present one) definitely suggest that this choice is reasonable.
- We expected that each region is coincident with one gene in average. Since the number of selected regions, 507, is almost equivalent to the number of gene symbols in these 507 regions, 525 (see supplementary materials), employment of region of 25,000 nucleic acid length seems to be reasonable.
- Since average gene length on human genome is ≥ 3 × 10^4^, the selection of region of 25,000 nucleic acid length is supposed to be association of one gene in each region. As denoted in the above, this expectation was fulfilled.

Although the above discussion might be enough to rationalize the usage of region of 25,000 nucleic acid length, we tried an alternative strategy as described in Materials and Methods. We downloaded peak call data set from GEO and tried to identify overlaps between peak regions. As a result, we could find only 22 regions of mean length of 5000 nucleic acid, with which only 13 gene symbols were associated. This tells us two things. Smaller region length, 5,000, results in regions without gene symbols. Shorter region length reduces the number of commonly identified regions between multiple experiments. This prevents us from performing downstream analysis. This failure of an alternative approach definitely suggests the suitability of the selection of region of 25,000 nucleic acid length.

One might also wonder if TFs can also work in cell line specific ways; thus there might be no reasons to select TFs common between two cell lines. It is really true that TFs can work in cell line specific ways; nevertheless, what we are interested in is a more robust bookmark that can likely work in mitoic process universally. If we selected TFs that work in cell line specific manner, it reduces the possibility that selected TFs work universally in mitoic process. The reason why we validated the selected genes based upon Enrichr that might include the results for other cell lines than U2OS and RPE1 is similar; if the selected genes are coincident with data bases retrieved from other cell lines, results are more unlikely accidental and are more likely robust and universal.

In this study, reliability of selected genes was evaluated by enrichment analysis. Since we have selected very small amount of genes, as small as c.a. 500, it is very unlikely for them to be associated with numerous enrichment. In spite of that, since our selected genes are associated with so many TF activities, we can assume that our selection of genes are reasonable. In the case that we cannot find any enrichment, we regard that our selection of singular value vectors is failure and we try to check if other selections can work better or not. It is worth noting that because other methods are not designed to deal with the studied problem, applying these methods generate inferior outcomes.

We show that selected TFs are expressive in cell lines as follows: First of all, we evaluated TFs by not only binding to genome but also co-occurrence with selected genes (e.g. (III) and (V) in Table 3). Thus, it is very likely that some TFs are expressive in cell lines where the selected genes are expressive. Second, we seek GEO Profiles in order to see if these TFs are expressive in U2OS cell lines and RPE cells. Then, we have found that almost all TFs were expressive in both U2OS cell lines and RPE cells in GEO profiles (see Additional file 4). Thus, it is not unreasonable to expect the expression of these TFs in two cell lines used in this study.

## 4 Conclusions

We applied a novel TD-based unsupervised FE method to various histone modifications across the whole human genome, and the levels of these modifications were measured during mitotic cell division to identify genes that are significantly associated with histone modifications. Potential bookmark TFs were identified by searching for TFs that target the selected genes. The TFs identified were functionally related to the cell division cycle, suggesting their potential as bookmark TFs that warrant further exploration.

## Author Contributions

YT planned and performed the study. YT and TT discussed the results and wrote the paper. All authors contributed to the article and approved the submitted version.

## Acknowledgments

This manuscript will be released as a pre-print at BioRxiv.

## Supplemental Data

Additional file 1: Genes identified by TD-based unsupervised FE; Additional file 2: Potential TFs that target identified genes (in Additional file 1) identified by Enrichr; Additional file 3: Sample R code used in the analyses performed in this study. Additional file 4: Expression of TFs selected in this study in GEO profiles.

## Data Availability Statement

All datasets analyzed in this study were obtained from GEO: GSE141139

