## Supplementary material for "Unsupervised tensor decomposition-based method to extract candidate transcription factors as histone modification bookmarks in post-mitotic transcriptional reactivation": Additional_file_3.docx

#batch.R to sum bw files within 25000 bps block

\# n is taken to be from 1 to 12 in order to process all 60 bw files

require(rtracklayer)

files <- unlist(strsplit(system("tar -tvf GSE141139_RAW.tar",intern=T)," "))

files <- files[grep("GSM",files)]

files <- files[grep(".bw",files)]

which <- GRanges("chr1", IRanges(1, 30000))

n=1

for (j in c((2+(n-1)*5):(2+n*5-1)))

{

system(paste("tar -xvf GSE141139_RAW.tar",files[j]))

dir <- gsub(".bw","",files[j])

system(paste("mkdir",dir))

bw <- import(files[j],which=which)

for (k in c(1:length(bw@seqinfo@seqnames)))

{

seqname <- bw@seqinfo@seqnames[k]

if (bw@seqinfo@seqlengths[k]>=25000)

{

breaks<-seq(1,bw@seqinfo@seqlengths[k],by=25000)

} else {

breaks<- c(1,bw@seqinfo@seqlengths[k])

}

Chip <- NULL

for (i in (c(1:(length(breaks)-1))))

{

cat(i," ")

which <- GRanges(seqname, IRanges(breaks[i], breaks[i+1]-1))

bw <- import(files[j],which=which)

Chip <- c(Chip,sum(bw$score))

}

save(file=paste(dir,"/Chip_",seqname,sep=""),Chip)

}

system(paste("rm",files[j]))

}

#----- prepare sample and dirs

require(readODS)

dirs <- list.files(pattern="GSM")

sample <- read.ods("sample.ods",sheet=1)

sample <- t(data.frame(strsplit(sample[,2],"-")))

sample <- data.frame(t(data.frame(strsplit(sample[,1]," "))),sample[,2:5])

rownames(sample) <- NULL

sample[,3] <- gsub("telopphase","telophase",sample[,3])

#------

#summing up 60 files into individual chromosome as tensor format

chr <- paste("chr",c(1:22,"M","X","Y"),sep="")

for (l in c(1:length(chr)))

{

cat(l," ")

x<-NULL

for (i in c(1:length(dirs)))

{

#cat (i, " ")

load(paste(dirs[i],"/Chip_",chr[l],sep=""))

x <- cbind(x,Chip)

}

cell <- names(table(sample[,1]))

histone <- names(table(sample[,2]))[-1]

phase <- names(table(sample[,3]))

rep <- names(table(sample[,6]))[1:2]

Z <- array(NA,c(dim(x)[1],2,4,3,2))

for (i in c(1:2)){

for (j in c(1:4)){

for (k in c(1:3)){

for (m in c(1:2)){

id <- intersect(intersect(intersect(grep(cell[i],sample[,1]),grep(histone[j],sample[,2])),

grep(phase[k],sample[,3])),grep(rep[m],sample[,6]))

Z[,i,j,k,m] <- x[,id]

}

}

}

}

save(file=paste("Z_",chr[l],sep=""),Z)

}

#-----

#bing all chtomosme files into one tensor

require(abind)

dnum <-NULL

Z_all <- NULL

for(l in c(1:length(chr)))

{

load(paste("./Z/Z_",chr[l],sep=""))

dnum<-c(dnum,dim(Z)[1])

Z_all <- abind(Z_all,Z,along=1)

}

#------ perform tensor decomposition --

require(rTensor)

Z_all <- apply(Z_all,2:5,scale)

HOSVD <- hosvd(as.tensor(Z_all)

#---- gene selection ----

P <- pchisq(scale(HOSVD$U[[1]][,4])^2,1,lower.tail=F)

require(biomaRt)

grch37 = useMart(biomart="ENSEMBL_MART_ENSEMBL", host="grch37.ensembl.org", path="/biomart/martservice", dataset="hsapiens_gene_ensembl")

gene_all <-NULL

for (l in c(1:22)){

cat("\n l= ",l,"\n")

if (l==1)

{

id <- c(1:dnum[l])[(p.adjust(P,"BH")<0.01)[1:dnum[l]]]

}else{

id <- c(1:dnum[l])[(p.adjust(P,"BH")<0.01)[sum(dnum[1:(l-1)])+c(1:dnum[l])]]

}

reg<-id*25000

reg<-data.frame(reg-24999,reg)

gene <- NULL

for (i in c(1:dim(reg)[1]))

{

cat(i," ")

gene_lst0 <- getBM(attributes=c('hgnc_symbol','entrezgene_id','chromosome_name','start_position','end_position'),filters = c('chromosome_name','start','end'), values=list(l,reg[i,1],reg[i,2]), mart = grch37)

gene <- rbind(gene,gene_lst0)

}

gene <- gene[!is.na(gene[,1]),]

gene <- gene[!is.na(gene[,1]) & gene[,1]!="",]

gene_all <- rbind(gene_all,gene)

}

write.table(file="gene_all.csv",gene_all,row.names=F,col.names=F,quote=F,sep="\t")

#DESeq2

#i= 1 and 4, j= 1 to 4

i<-2;j<-4

Z1 <- Z_all[,i,j,,]

Z1 <- data.frame(Z1[,1,],Z1[,2,],Z1[,3,])

colnames(Z1) <- 1:6

coldata <- data.frame(rep(1:3,each=2))

colnames(coldata) <- "condition"

coldata$condition <- factor(coldata$condition)

Z1 <- trunc(Z1)

dds <- DESeqDataSetFromMatrix(countData = Z1,colData = coldata,design = ~ condition)

featureData <- data.frame(gene=rownames(Z1))

mcols(dds) <- DataFrame(mcols(dds), featureData)

mcols(dds)

dds <- DESeq(dds)

res <- results(dds)

table(res$padj<0.01)

#csaw

#--- mapping fastq and generate sorted bam--

\# should be execued to 60 fastq files outside of R

nohup ~/sratoolkit.2.9.2-centos_linux64/bin/fastq-dump SRR10540100 --gzip > nohup_SRR10540100.out &

~/bowtie2-2.3.5.1-linux-x86_64/bowtie2 -x ~/Homo_sapiens/UCSC/hg38/Sequence/Bowtie2Index/genome SRR10540100.fastq.gz -p 4 -S SRR10540100.sam 2> nohup_SRR10540100.out &

nohup samtools view -Sb SRR10540100.sam | samtools sort - -o SRR10540100_sorted.bam > nohup_SRR10540100_sort.out

samtool index SRR10540100_sorted.bam

#-----

bam.files <- list.files("./",pattern="bam",recur=T)

bam.files <- bam.files[-grep("bai",bam.files)]

#bam.files <- bam.files[1:6] #U2OS H3K4me3

#bam.files <- bam.files[13:18] #U2OS H3K4me1

#bam.files <- bam.files[34:39] #RPE1 H3K4me3

#bam.files <- bam.files[46:51] #RPE1 H3K4me1

#bam.files <- bam.files[7:12] #U2OS H3K27ac

#bam.files <- bam.files[40:45] #RPE1 H3K27ac

#bam.files <- bam.files[c(25,28,26,29,27,30)] #U2OS Input

#bam.files <- bam.files[c(52,55,53,56,54,57)] #RPE1 Input

library(csaw)

require(edgeR)

param <- readParam(minq=20)

data <- windowCounts(bam.files, ext=110, width=10, param=param)

keep <- aveLogCPM(asDGEList(data)) >= -1

filtered.data <- data[keep,]

binned <- windowCounts(bam.files, bin=TRUE, width=10000, param=param)

data <- normFactors(binned, se.out=data)

y <- asDGEList(data)

cell.type<-c(rep(1:3,each=2))

design <- model.matrix(~factor(cell.type))

y <- estimateDisp(y, design)

summary(y$trended.dispersion)

require(statmod)

fit <- glmQLFit(y, design, robust=TRUE)

summary(fit$var.post)

results <- glmQLFTest(fit)

head(results$table)

table(p.adjust(results$table[,4],"BH")<0.01)
